# Psychedelics and schizophrenia: Distinct alterations to Bayesian inference

**DOI:** 10.1101/2022.01.31.478484

**Authors:** Hardik Rajpal, Pedro A.M. Mediano, Fernando E. Rosas, Christopher B. Timmermann, Stefan Brugger, Suresh Muthukumaraswamy, Anil K. Seth, Daniel Bor, Robin L. Carhart-Harris, Henrik J. Jensen

## Abstract

Schizophrenia and states induced by certain psychotomimetic drugs may share some physiological and phenomenological properties, but they differ in fundamental ways: one is a crippling chronic mental disease, while the others are temporary, pharmacologically-induced states presently being explored as treatments for mental illnesses. Building towards a deeper understanding of these different alterations of normal consciousness, here we compare the changes in neural dynamics induced by LSD and ketamine (in healthy volunteers) against those associated with schizophrenia, as observed in resting-state M/EEG recordings. While both conditions exhibit increased neural signal diversity, our findings reveal that this is accompanied by an increased transfer entropy from the front to the back of the brain in schizophrenia, versus an overall reduction under the two drugs. Furthermore, we show that these effects can be reproduced via different alterations of standard Bayesian inference applied on a computational model based on the predictive processing framework. In particular, the effects observed under the drugs are modelled as a reduction of the precision of the priors, while the effects of schizophrenia correspond to an increased precision of sensory information. These findings shed new light on the similarities and differences between schizophrenia and two psychotomimetic drug states, and have potential implications for the study of consciousness and future mental health treatments.

## 1. Introduction

Classic serotonergic psychedelic drugs have seen a blooming resurgence among the public and the scientific community in recent years, largely driven by promising clinical research into their therapeutic potential [1, 2]. At the same time, and somewhat paradoxically, psychedelics are known to elicit effects that mimic some symptoms of psychosis-earning them the label of ‘psychotomimetic drugs’ [3]. Elucidating the similarities and differences between the psychedelic state and schizophrenia is an important neuroscientific challenge that could deepen our understanding of the substantive alterations of perception, cognition, and conscious experience induced by these conditions, while paving the way towards safe and effective psychedelic therapy.

To contrast these conditions in an empirical manner, we compare neuroimaging data from patients suffering from schizophrenia and healthy subjects under the effects of two psychoactive substances: the classical psychedelic lysergic acid diethylamide (LSD) [4] and the dissociative drug ketamine (KET) [5]. Using standardised assessments, it has been claimed that KET reproduces both positive and negative symptoms of schizophrenia in humans [6], and its mechanism of action – NMDA receptor antagonism – is thought to reproduce a key element of the molecular pathophysiology of schizophrenia [7, 8]. LSD – in common with all classical psychedelics – is a potent agonist of a number of serotonin receptors, but its characteristic effects depend primarily on 5-HT_2A_ [9]. These neurotransmitter systems are linked to psychosis (in particular visual perceptual changes), and are implicated in psychotic disorders like schizophrenia – although see Ref. [10] for an alternative account.

Both psychotomimetic drug states and schizophrenia are also associated with marked changes in large-scale neural dynamics. For both LSD and KET, previous studies have found increased signal diversity in subjects’ neural dynamics [11, 12] and reduced information transfer between brain regions [13]. However, in the case of KET, evidence from intracranial recordings in cats suggests a much more complicated picture than that of LSD, with very high variability across individuals, brain regions, and dose levels [14]. In a separate line of enquiry, work on EEG data from patients with schizophrenia has also found increased signal diversity [15, 16], akin to the effect found under these drugs. Nonetheless, a parsimonious account explaining the similarities and differences between the two states is still lacking.

A promising approach to gain insights into the mechanisms driving the core similarities and differences between psychotomimetic drug states and schizophrenia is to leverage principles from the predictive processing (PP) framework of brain function [17, 18]. A key postulate of the PP framework is that the dynamics of neural populations can be viewed as engaged in processes of inference involving top-down and bottom-up signals. Under this framework, brain activity can be viewed as resulting from a continuous modelling process in which a prior distribution interacts with new observations via incoming sensory information. In accordance with principles of Bayesian inference, discrepancies between the prior distribution and incoming signals (called ‘prediction errors’) carried by the bottom-up signals drive revisions to the top-down activity, so as to minimize future surprise.

The PP framework has been used to explain perceptual alterations observed in both psychotomimetic drug states [19, 20] as well as in psychiatric illnesses [21] with a focus on schizophrenia [22, 23, 24, 25]. PP has also been used to understand the action of psychedelics, most notably through the ‘relaxed beliefs under psychedelics’ (or REBUS) model [26] which posits that psychedelics reduce the precision of prior beliefs encoded in spontaneous brain’s activity. REBUS has also been used to inform thinking on the therapeutic mechanisms of psychedelics, where symptomatology can be viewed as pathologically over-weighted beliefs or assumptions encoded in the precision weighting of brain activity encoding them.

To deepen our understanding of the similarities and differences between these conditions, in this paper we replicate and extend findings on neural diversity and information transfer under the two psychotomimetic drugs (LSD and KET) and in schizophrenia using EEG and MEG recordings, and we reproduce these experimental findings as perturbations to a single PP model. Our modelling results reveal that the effects observed under the drugs are indeed reproduced by decreasing the precision-weighting of the priors, while the effects observed under schizophrenia are reproduced by increased precision-weighting of the bottom-up sensory information. Overall, this study puts forward a more nuanced understanding of the relationship between two different psychotomimetic drug states and schizophrenia, and offers a new model-based perspective on how these conditions alter conscious experience.

## 2. Materials and Methods

### 2.1. Data acquisition and preprocessing

Data from 29 patients diagnosed with schizophrenia and 38 age-matched healthy control subjects were obtained from the Bipolar-Schizophrenia Network on Intermediate Phenotypes (BSNIP) database [27]. The subjects were selected within an age range of 20-40 years to match the psychedelic datasets described below. Data included 64-channel EEG recordings sampled at 1000 Hz of each subject in eyes-closed resting state, along with metadata about demographics (age and gender), and patients’ medications. The strength of the medication was estimated using the number of antipsychotics taken by each patient (mean: 2.7, range: 0-8), as the dosage of each medication was not available.

Data from healthy subjects under the effects of both drugs was obtained from previous studies with LSD [4] (N = 17) and ketamine [28] (N = 19). Data included MEG recordings from a CTF 275-channel axial gradiometer system with a sampling frequency of 600 Hz. Each subject underwent two scanning sessions in eyes-closed resting state: one after drug administration and another after a placebo (PLA).

Preprocessing steps for all datasets were kept as consistent as possible, and were performed using the Fieldtrip [29] and EEGLAB [30] libraries. First, the data was segmented into epochs of 2 seconds, and epochs with strong artefacts were removed via visual inspection. Next, muscle and eye movement artefacts were removed using ICA [31]. Then, a LCMV beamformer [32] was used to reconstruct activity of sources located at the centroids of regions in the Automated Anatomical Labelling (AAL) brain atlas [33]. Finally, source-level data was bandpass-filtered between 1– 100 Hz, and downsampled with phase correction to 250 Hz (EEG) and 300 Hz (MEG), and AAL areas were grouped into 5 major Regions of Interest (ROIs): frontal, parietal, occipital, temporal and sensorimotor (see Figure 1 and Table B.1 in the Appendix). In the rest of the paper we refer to these 5 areas as “ROIs” and to the AAL regions as “sources.”

**Figure 1:**
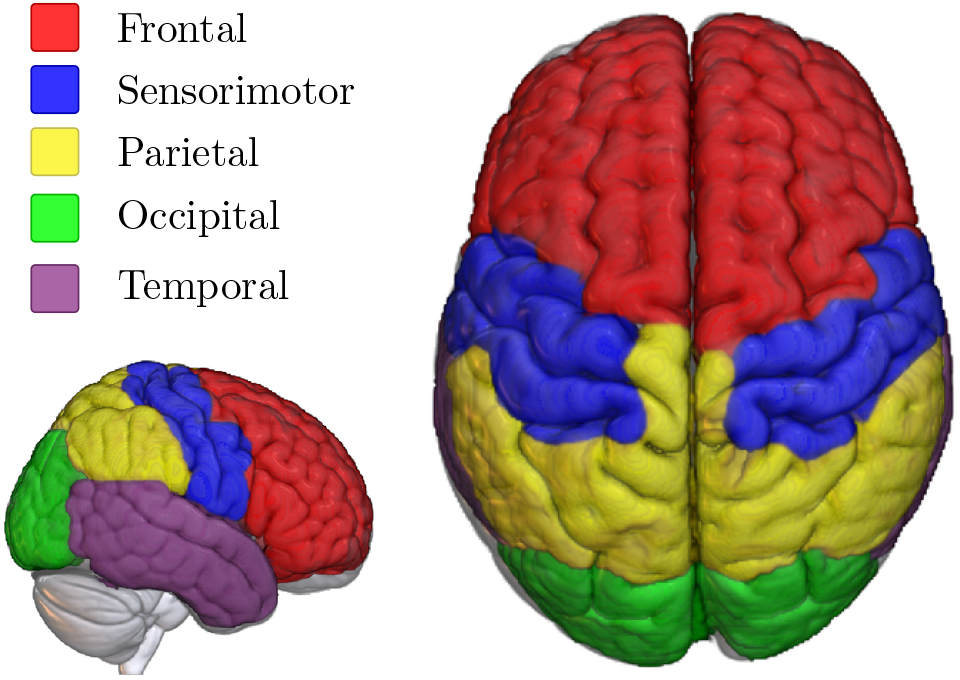
Regions of interest (ROIs) represented on the MNI-152 standard template. Each ROI is comprised of several regions of the AAL atlas, as per Table B.1.

### 2.2. Analysis metrics

Our analyses are focused on two complementary metrics of neural activity: Lempel-Ziv complexity (LZ) and transfer entropy (TE). Both metrics are widely used and robustly validated measures of neural dynamics across states of consciousness [13, 34, 11, 12].

Lempel-Ziv complexity (LZ) is a measure of the diversity of patterns observed in a discrete – typically binary – sequence. When applied to neuroimaging data, lower LZ (with respect to wakeful rest) has been associated with unconscious states such as sleep [35] or anaesthesia [36], and higher LZ with states of richer phenomenal content under psychedelics, ketamine [11, 12] and states of flow during musical improvisation [37].

To calculate LZ, first one needs to transform a given signal of length *T* into a binary sequence. For a given epoch of univariate M/EEG data, we do this by calculating the mean value and transforming each data point above the mean to 1 and each point below to 0. Then, the resulting binary sequence is scanned sequentially using the LZ76 algorithm presented by Kaspar and Schuster [38], which counts the number of distinct “patterns” in the signal. Finally, following results by Ziv [39], the number of patterns is divided by log_2_(*T*)/*T* to yield an estimate of the signal’s entropy rate [40], which we refer to generically as LZ. This process is applied separately to each source time series (i.e. to each AAL region), and the resulting values are averaged according to the grouping in Table B.1 to yield an average LZ value per ROI.

In addition to LZ, our analyses also consider transfer entropy (TE) [34] — an information-theoretic version of Granger causality [41] — to assess the dynamical interdependencies between ROIs. The TE from a source region to a target region quantifies how much better one can predict the activity of the target after the activity of the source is known. This provides a notion of directed functional connectivity, which can be used to analyse the structure of large-scale brain activity [13, 42].

Mathematically, TE is defined as follows. Denote the activity of two given ROIs at time *t* by the vectors ***X***_*t*_ and ***Y***_*t*_, and the activity of the rest of the brain by ***Z***_*t*_. Note that ***X***_*t*_, ***Y***_*t*_, and ***Z***_*t*_ have one component for each AAL source in the corresponding ROI(s). TE is computed in terms of Shannon’s mutual information, *I*, as the information about the future state of the target, ***Y***_*t*+1_, provided by ***X***_*t*_ over and above the information in ***Y***_*t*_ and ***Z***_*t*_:

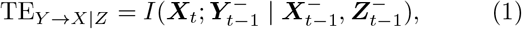

where 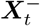 refers to the (possibly infinite) past of ***X***_*t*_, up to and including time *t* (and analogously for ***Y***_*t*_ and ***Z***_*t*_). This quantity can be accurately estimated using state-space models with Gaussian innovations [43], implemented using the MVGC toolbox [44]. Note that, when calculating the TE between ROIs, we consider each ROI as a vector — without averaging the multiple AAL sources into a single number. The result is a directed 5 × 5 network of conditional TE values between pairs of ROIs, which can be tested for statistical differences across groups.

### 2.3. Statistical analysis

For both LSD and KET datasets, since the same subjects were monitored under both drug and placebo conditions, average subject-level differences (either in LZ or TE) were calculated for each subject, and one-sample t-tests were used on those differences to estimate the effect of the drug.

For the data of patients and controls in the schizophrenia dataset, group-level differences were estimated via linear models. These models used either LZ or TE as target variable, and condition (schizophrenia or healthy), age, gender, and number of antipsychotics (set to zero for healthy controls) as predictors. Motivated by previous work suggesting a quadratic relationship between complexity and age [45], each model was built with either a linear or quadratic dependence on age, and the quadratic model was selected if it was preferred over a linear model by a log-likelihood ratio test (with a critical level of 0.05).

Finally, multiple comparisons when comparing TE values across all pairs of ROIs were addressed by using the Network-Based Statistic (NBS) [46] method, which identifies ‘clusters’ of differences – i.e. connected components where a particular null hypothesis is consistently rejected while controlling for family-wise error rate. Our analysis used an in-house adapted version of NBS that works on directed networks, such as the ones provided by TE analyses.

### 2.4. Computational modelling

A computational model was developed in order to interpret the LZ and TE findings observed on the neuroimaging data. Building on *predictive processing* principles [18], we constructed a Bayesian state-space model that provides an idealised common ground to contrast the three studied conditions – the psychotomimetic drug states, schizophrenia, and baseline (i.e. healthy controls). Our modelling is based on the postulate that the activity of neuronal populations across the brain can be interpreted as carrying out inference on the causes of their afferent signals. Following this view, the proposed model considers the following elements:

- the internal state of a low-level region (i.e. near the sensory periphery), denoted by *s*_*t*_;
- the internal state of neural activity taking place functionally one level above, denoted by *h*_*t*_;
- the signal generated at the high-level region in the form of a prediction of the low-level activity, denoted by *ŝ*_*t*_;
- the signal generated at the low-level region in the form of a prediction error *ξ*_*t*_; and
- the precision of the prior *λ*_p_ and precision of sensory/afferent information *λ*_s_.

This model represents neural activity within a larger hierarchical processing structure, as illustrated in Figure 2. The key principle motivating this model is that minimisation of prediction error signals throughout the hierarchy, by updating top-down predictions, implements a tractable approximation to Bayesian inference.^2^

**Figure 2:**
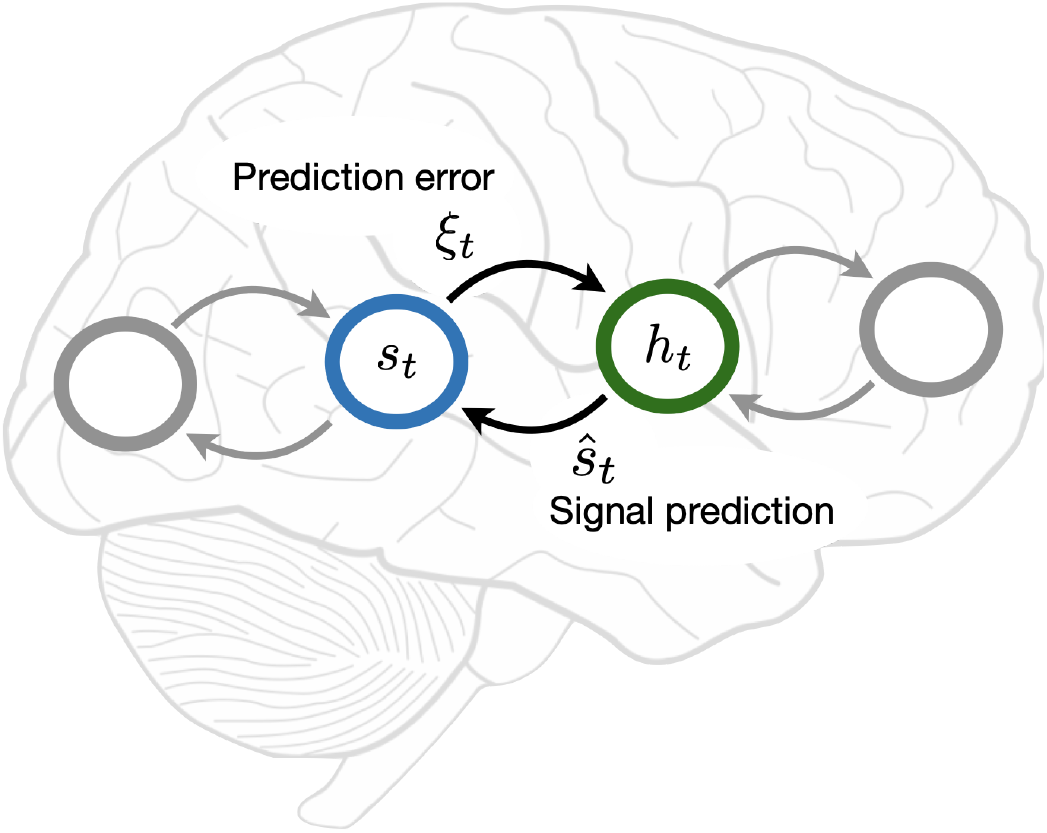
Graphical illustration of the predictive processing model. The activity of a high-level neural population is represented as a prediction *ŝ*_*t*_, and the activity of a low-level population as a prediction error *ξ*_*t*_. The internal states of the high- and low-level regions are captured by *s*_*t*_ and *h*_*t*_, respectively.

Within this model, we represent the schizophrenia and psychedelic conditions as different types of disruption to Bayesian inference. To describe the psychedelic state, we build on the REBUS hypothesis [26], which posits a reduced precision-weighting of prior beliefs, leading to increased bottom-up influence. Conversely, to describe schizophrenia we build on the canonical predictive processing account of psychosis in schizophrenia [48], which postulates an increased precision of sensory input, along with decreased precision of prior beliefs [22, 21]. Therefore, both conditions are similar in that there is a relative strengthening of bottom-up influence, although instantiated in different ways – which, as shown in Sec. 3.3, bears important consequences for the behaviour of the model.

It is important to note that predictive processing accounts of schizophrenia remain hotly debated, with other works proposing an increase of prior precision (instead of decrease) as a model of auditory and visual hallucinations [49, 50]. Recent reviews [48] have attempted to reconcile both views by suggesting that sensory hallucinations may be caused by stronger priors, while hallucinations related to self-generated phenomena (like inner speech or self-attention [51]) may stem from weaker priors. Here, we base our modelling of SCZ on the weak prior hypothesis, as described above - we return to this issue in this discussion.

To simulate the aberrant dynamics of the inference process, as described above, we consider a given afferent signal (*s*_*t*_) and construct the corresponding activity of a higher area (*h*_*t*_), prediction (*ŝ*_*t*_), and prediction error (*ξ*_*t*_), building on the rich literature of state-space models in neuroscience [52, 53, 54]. Specifically, we use the linear stochastic process:

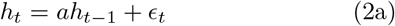

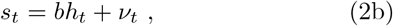

where *a, b* are weights, and *ϵ*_*t*_, *ν*_*t*_ are zero-mean Gaussian terms with precision (i.e. inverse variance) *λ*_p_ and *λ*_s_, respectively. Note that this formulation is equivalent to

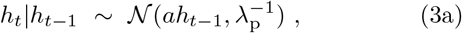

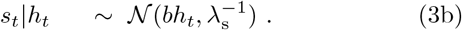

As we show in the following, *λ*_p_ corresponds to the precision of the prior and *λ*_s_ to the precision of sensory/afferent information.

The dynamics of this system can be described as a recurrent update between predictions and prediction errors as follows. Eq. (3b) implies that the internal state *h*_*t*_ generates a prediction about the low-level activity given by *ŝ*_*t*_ = *bh*_*t*_. At the same time, the dynamics of the high-level region can be seen as a Bayesian update of *h*_*t*_ given *s*_*t*_ and *h*_*t*−1_. Under some simplifying assumptions, the mean of the posterior distribution of *h*_*t*+1_ (denoted by *ĥ*_*t*+1_) is equal to (see Appendix A)

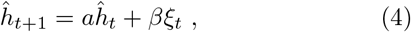

which effectively combines a *prior aĥ*_*t*_ (which is the optimal prediction of *h*_*t*+1_ given only *ĥ*_*t*_, as seen from Eq. (3a)) and a *likelihood* given by the prediction error *ξ*_*t*_ = *s*_*t*_ − *bĥ*_*t*_ that is precision-weighted via *β*, a parameter known as the *Kalman gain* [55].

In our simulations, the model is first calibrated using as afferent signals (i.e. *s*_*t*_) data from the primary visual cortex, corresponding to epochs randomly sampled from the placebo conditions in the LSD and KET datasets. This calibration results in the estimation of the model parameters *a*^*con*^, *b*^*con*^,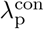,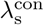 for the control condition, which is done using the well-known expectation-maximisation algorithm [56]. With these, the schizophrenia condition is then modelled by setting

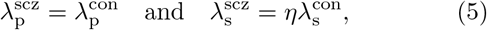

where *η* > 1 is referred to as a *noise factor*. This increase of *λ*_*s*_ induces a strengthening of bottom-up prediction errors, and makes the posterior of *h*_*t*_ excessively precise. Conversely, the drug condition is modelled by setting

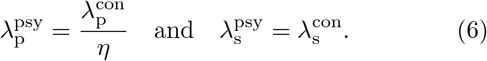

Reducing *λ*_p_ also increases the influence of prediction errors, but reduces the precision of the posterior of *h*_*t*_. Subsequently, for both conditions the parameters *a, b* are retrained with another pass of the expectation-maximisation algorithm on the placebo trials.

Finally, to compare the model with the empirical M/EEG data, the LZ of the neural activity elicited in the low-level area (i.e. the prediction errors, *ξ*_*t*_) and the top-down transfer entropy (from the high-level activity *ŝ*_*t*_ towards the low-level activity *ξ*_*t*_) are calculated for each of these three models (control, schizophrenia, and drug).

## 3. Results

### 3.1. LSD, KET and schizophrenia all show increased LZ

We begin the analysis by comparing changes in signal diversity, as measured by LZ, across the LSD, ketamine (KET), and schizophrenia (SCZ) datasets.

Our results show strong and significant increases in LZ in all three datasets (Fig. 3), in line with previous work [11, 12, 15, 16]. In all three cases the LZ increases are widespread throughout the brain, with the effects in schizophrenia patients being more pronounced in frontal and parietal regions. While the t-scores are higher in LSD and KET than schizophrenia, this could be due to the within-subjects design of both drug experiments – which are more statistically powerful than the between-subjects analysis used on the schizophrenia dataset.

**Figure 3:**
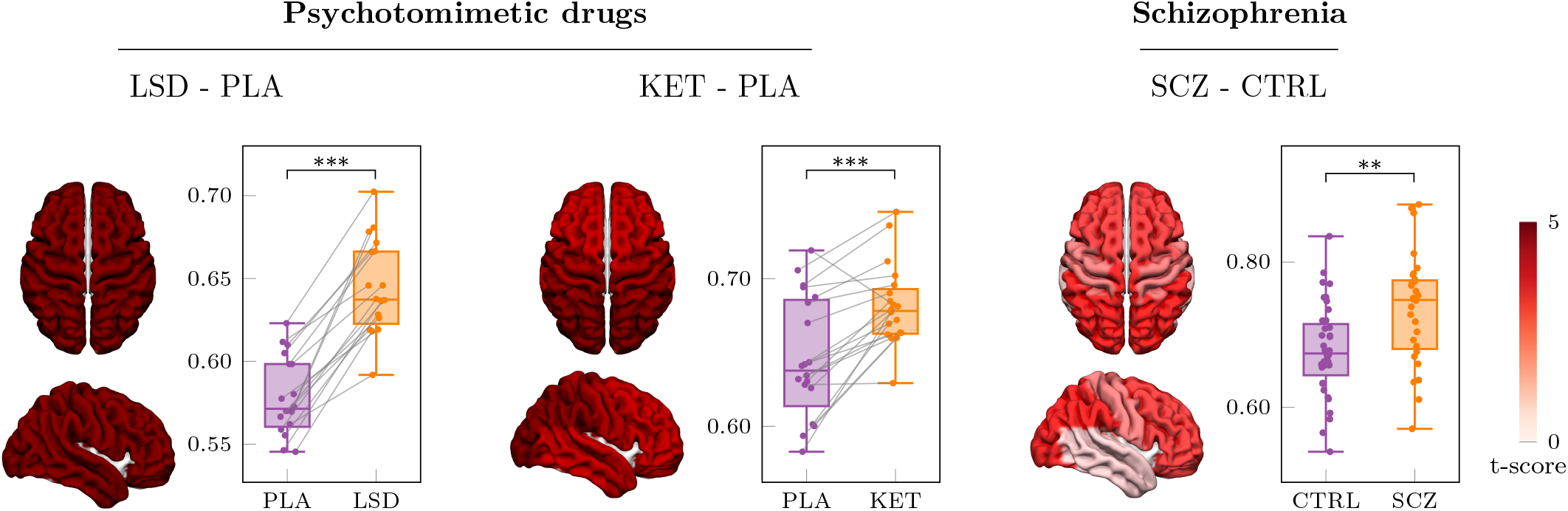
Increased signal diversity in subjects under the effects of psychotomimetic drugs and in schizophrenia patients. LZ changes are widespread across all ROIs in the three datasets. For the schizophrenia dataset, LZ values shown are corrected for age, gender, and the number of antipsychotic medications taken by each patient using a linear model.

Interestingly, we found that controlling for the medication status of each schizophrenia patient was crucial to obtain results that match prior work [15]. A direct comparison of LZ values between patients and controls yielded no significant differences (*t* = −0.38, *p* = 0.70); however, when using a linear model correcting for age, gender, and number of antipsychotics, the antipsychotics coefficient of the model reveals a negative effect on LZ (*β* = −0.016, *t* = −2.3, *p* = 0.021). Additionally, a two-sample t-test calculated between the corrected LZ values of patients and controls yields a substantial difference (*t* = 3.4, *p* = 0.001). Nonetheless, the sensitivity of this result to these pre-processing steps means it should be considered preliminary and explored further in future research (see the corresponding discussion in Sec. 4.3).

### 3.2. Opposite effect of psychotomimetic drugs and schizophrenia on information transfer

We next report the effects of LSD, KET, and schizophrenia on large-scale information flow in the brain, as measured via transfer entropy (TE). The TE between each pair of ROIs (conditioned on all other ROIs) is calculated for each subject, and used to build directed TE networks. The resulting networks were tested for differences between the drug states and placebo conditions (for LSD and KET), and between patients and controls (for SCZ), correcting for multiple comparisons via cluster permutation testing (see Sec. 2.3).

We found a ubiquitous decrease in the TE between most pairs of ROIs under LSD and KET (Fig. 4), which is consistent with previous findings [13]. In contrast, SCZ patients exhibit marked localised increases in TE – and no decreases – with respect to the control subjects. Notably, most increases in TE originated in the frontal ROI, and are strongest between the frontal and occipital ROIs. The increase of information transfer seen in schizophrenia patients therefore takes place “front to back” – aligned with the pathways thought to carry top-down information in the brain from highly cognitive, decision-making regions to unimodal regions closer to the sensory periphery.

**Figure 4:**
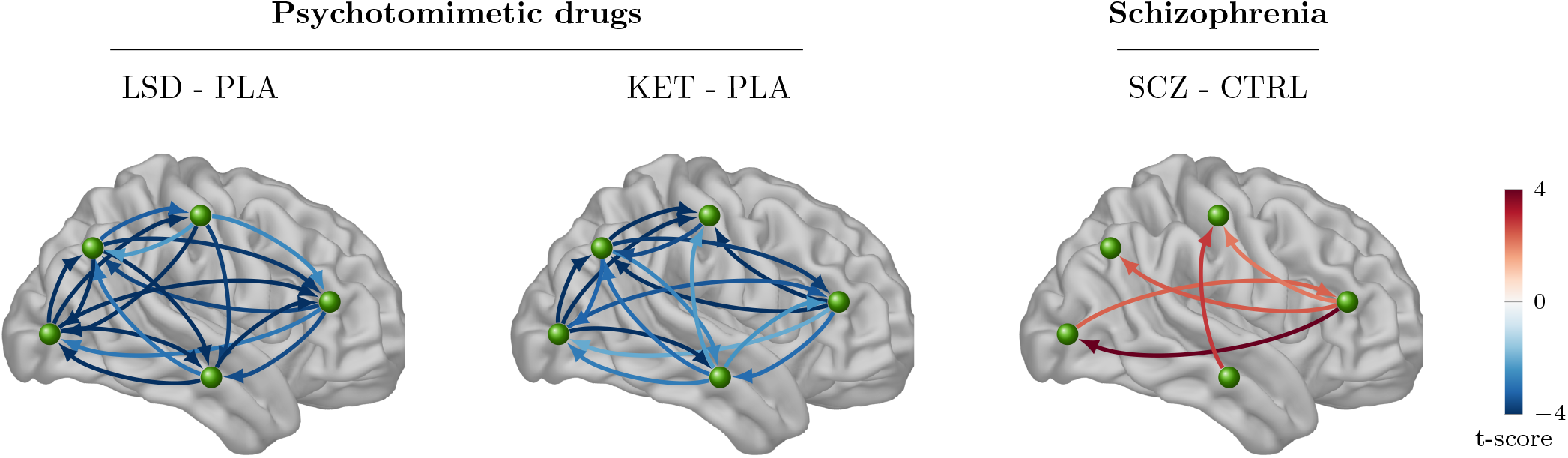
Lower information transfer under LSD and ketamine but higher information transfer for schizophrenia patients. Transfer entropy (TE) shows a strong widespread decrease in subjects under the effect of LSD or ketamine (KET), compared to a placebo. Conversely, schizophrenia (SCZ) patients show an increase in TE with respect to controls (CTRL), especially from the frontal region to the rest of the brain (controlling for age, gender and antipsychotic use). Links shown are significant after multiple comparisons correction.

As was the case for LZ, controlling for antipsychotic use was key to revealing differences between the healthy controls and schizophrenia patients. In addition, we found a small negative correlation between antipsychotic use and TE between certain ROI pairs – but, unlike for LZ, this effect did not survive correction for multiple comparisons.

### 3.3. Computational model reproduces experimental results

So far, we have seen that subjects under the effects of two different psychotomimetic drugs display increased signal diversity and reduced information flow in their neural dynamics. In comparison, schizophrenia patients display increased complexity but also increased information flow with respect to healthy controls. We now show how complementary perturbations to the precision terms of the predictive processing model introduced in Section 2.4 reproduce these findings.

We compared the basline model against the drug and schizophrenia variants by systematically increasing the noise factor *η*, which results in reduced prior precision in the drug model, and increased sensory precision in the schizophrenia model. We then computed the corresponding LZ and TE based on the model-generated time series *ξ*_*t*_, *ŝ*_*t*_ as per Sec. 2.4 (Fig. 5).

**Figure 5:**
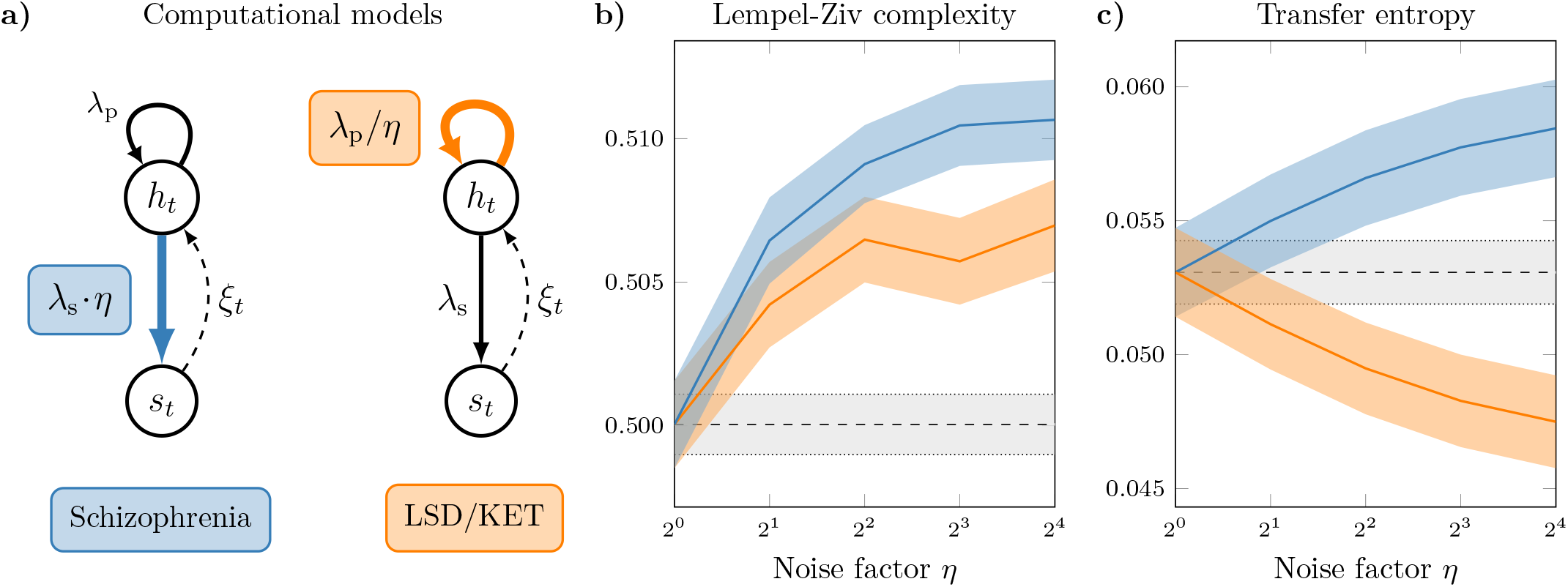
A computational model based on predictive processing principles reproduces experimental findings in the LSD, ketamine and schizophrenia datasets. (**a**) By increasing the sensory precision (for schizophrenia; **blue**), or reducing the prior’s precision (for LSD and KET; **orange**) by a given ‘noise’ factor *η*, the model can reproduce the experimental findings of (**b**) increased in LZ in both conditions, and (**c**) opposite changes in TE in both conditions, compared to a baseline (grey).

Results show that the proposed model successfully reproduced the experimental findings of both LZ and TE under the two different psychotomimetic drugs and schizophrenia (Fig. 5). Interestingly, the model also shows (Fig. 5b) that a relative strengthening of sensory information (via either increased sensory precision, or decreased prior precision) can trigger either an increase or a decrease (respectively) of top-down transfer entropy. This suggests that transfer entropy changes cannot be directly interpreted as revealing the changes in any underlying predictive processing mechanisms (see Discussion).

Finally, as a control, we repeated the analysis on the model but exploring the variation of the precision terms in the two unexplored directions — either reducing *λ*_p_ or increasing *λ*_s_ (see Section 2.4). Neither of these changes reproduced the experimental findings (results not shown), which highlights the specificity of the modelling choices.

## 4. Discussion

In this paper we have analysed MEG data from healthy subjects under the effects of the psychotomimetic drugs LSD and ketamine, as well as EEG data from a cohort of schizophrenia patients and healthy control subjects. We focused on signal diversity and information transfer, both widely utilised metrics of neural dynamics. We found that all datasets show increases in signal diversity, but diverging changes in information transfer, which was higher in schizophrenia patients but lower for subjects under the effects of either drug. In addition to replicating previous results reporting signal diversity and information transfer under the effects of both drugs [11, 12, 13], we described new findings applying these metrics to schizophrenia.

Using a computational model inspired by predictive processing principles [18, 57], we showed that this combination of effects can be reproduced via specific alterations to prediction updating, which can be interpreted as specific forms of disruption to Bayesian inference. Critically, the effects of both psychotomimetic drugs and schizophrenia, on both signal diversity and information transfer, are explained by a relative strengthening of sensory information over prior beliefs, although triggered by different mechanisms – a decrease in the precision of priors in the case of psychotomimetic drugs (consistent with Ref. [26]), and an increase in the precision of sensory information for schizophrenia.

### 4.1. Increased sensory precision in schizophrenia

The idea that the symptoms of schizophrenia can be understood as alterations to processes of Bayesian inference has been particularly fertile in the field of computational psychiatry [21]. In particular, various studies based on PP have related psychosis to decreased precision of prior beliefs and increased precision of the sensory inputs [58, 19, 22, 59, 60]. These computational models have been supported by a growing number of related experimental findings, including an enhanced confirmation bias [61], impaired reversal learning [62, 63], and a greater resistance to visual illusions [64]. For instance, schizophrenia patients are less susceptible to the Ebbinghaus illusion, which arises primarily from misleading prior expectations, suggesting that patients do not integrate this prior context with sensory evidence and thus achieve more accurate judgements [65].

Most of the above mentioned studies are task-based, focusing on differentiating perceptual learning behaviours between healthy controls and schizophrenia patients. Though these studies provide a range of experimental markers, the corresponding methodologies cannot be applied to resting-state or task-free conditions, under which it is known that certain behavioural alterations (e.g. delusions, anhedonia, and paranoia) persist [66, 67].

The findings presented in this paper provide a step towards bridging this important knowledge gap by providing empirical and theoretical insights into resting-state neural activity under schizophrenia. Although we build on and replicate results related to signal diversity, we are not aware of previous studies of information transfer on schizophrenia in resting state.

### 4.2. Beyond unidimensional accounts of top-down vs bottom-up processing

The findings presented here link spontaneous brain activity to the PP framework using empirical metrics of signal diversity and information transfer. In the psychotomimetic drug condition, the former increases while the latter decreases; in schizophrenia, both increase – in both cases as compared to baseline placebo or control. The explanation for this pattern of results, articulated by our computational model, is based on the idea that a bias favouring bottom-up over top-down processing can be triggered by changing different precision parameters, which can give rise to opposite effects in specific aspects of the neural dynamics. This observation, we argue, opens the door to more nuanced analyses for future studies.

The increased transfer entropy from frontal to posterior brain areas observed under schizophrenia could be naively interpreted as supporting increased top-down regulation; however, neither the empirical analysis nor the computational model warrant this conclusion. Transfer entropy simply indicates information flow and is agnostic about cusing functional role. Moreover, our model-based analyses illustrate how aberrant Bayesian inference in which bottom-up influences become strong can trigger either an increase or a decrease in transfer entropy from frontal to posterior regions, depending on which precision terms are involved. An interesting possible explanation for this divergence between mechanisms and TE is provided by recent results that show that TE is an aggregate of qualitatively different *information modes* [68]. Future work may explore if resolving TE into its finer constituents might provide a more informative mapping from observed patterns to underlying mechanisms.

Taken together, these findings suggest that conceiving the bottom-up vs top-down dichotomy as a single-5 dimensional trade-off might be too simplistic, and that multi-dimensional approaches could shed more light on this issue. In particular, our results show how such a simplistic view fails to account for the rich interplay of similarities and differences between schizophrenia and psychosis.

### 4.3. Limitations and future work

While our empirical and modelling results agree with the canonical PP account of psychosis [48], some reports have suggested a stronger influence of priors over sensory signals especially in some cases of hallucinations [69, 70, 71]. It is important to remark that the ‘strengthened prior’ interpretation put forward by these task-based studies cannot be accounted for by the simple computational modelling developed here. Future work on multi-level extensions of this model could test if a richer hierarchical model could implicate a stronger influence of certain high-level priors under certain conditions.

Regarding the empirical analyses, it is important to note that our analyses are subject to a few limitations due to the nature of the data used. First, the analyses used only 60 AAL sources across 5 ROIs (due to the spatial resolution limitations of EEG), and therefore may neglect potential PP effects that may exist at smaller spatial scales. In addition, the studies on both drugs and schizophrenia used different imaging methods (MEG vs EEG), sampling rate, and experiment designs (within vs between subjects), complicating direct comparisons. Finally, future work should examine how power spectra across the different conditions relate to the findings presented, in terms of both their effect on LZ [72], and their relationship with top-down and bottom-up signalling, for example using band-limited Granger causality [73], as well as how directed functional connectivity measures relate to undirected measures such as mutual information and coherence [13].

Similarly, while the measures discussed here capture significant differences between schizophrenia patients and healthy controls, more work needs to be done to further characterise the differences within the schizophrenia spectrum, which features a heterogenous array of symptoms and states, e.g. at different phases of the so-called ‘psychotic process’ [74]. This might entail a closer look at the clinical symptom scores of the patients and their relationship to neural dynamics, which was not possible here due to the lack of appropriate metadata. An interesting possibility is that the neural underpinnings of positive and negative symptoms could be different [22], and investigating these differences may yield further insight into schizophrenia itself and its relationship with the psychotomimetic drug states. Moreover, both schizophrenia and drug-induced states can be conceived of as dynamic states of consciousness, comprised of several sub-states and/or episodes with hallucinations, delusions and negative symptoms varying widely between and within individuals. Future studies could explore these finer fluctuations in conscious state [75], as well as what features or episodes overlap in the neural and psychological levels between psychotomimetic drug states and schizophrenia.

Finally, recall that (as described in Sec. 2.1) we used the number of antipsychotic medications being used by each patient as a proxy measure for their medication load. We acknowledge that this is an oversimplification, and that richer datasets may allow a more detailed inspection of the effects of each particular medication – which would potentially bring more nuance to these analyses. Also, the models used for statistical analysis (as per Sec. 2.3) are linear and may not capture possible non-linear dependencies between antipsychotic use and its effect on neural dynamics (in our case, LZ or TE). Bearing this caveat in mind, the preliminary results regarding antipsychotic use suggest that they bring the patients’ neural dynamics closer to the range of healthy controls. This finding should be replicated with more detailed analyses, and, if robust, could potentially be used to investigate the mechanism of action of current antipsychotic drugs.

### 4.4. Final remarks

In this paper we have contrasted changes in brain activity in individuals with schizophrenia (compared to healthy controls) with changes induced by a classic 5-HT_2A_ receptor agonist psychedelic, LSD, and an NMDA antagonist dissociative, ketamine (compared to placebo). Empirical analyses revealed that both schizophrenia and drug states show an increase in neural signal diversity, but they have divergent transfer entropy profiles. Furthermore, we proposed a simple computational model based on the predictive processing framework [18] that recapitulates the empirical findings through distinct alterations to optimal Bayesian inference. In doing so, we argued that both schizophrenia and psychotomimetic drugs can be described as inducing a stronger “bottom-up” influence of sensory information, but in qualitatively different ways, thus painting a more nuanced picture of the functional dynamics of predictive processing systems. Crucially, the proposed model differs from others in the literature in that it is a model of resting-state (as opposed to task-based) brain activity, bringing this methodology closer to other approaches to neuroimaging data analysis based on complexity science [76].

Overall, this study illustrates the benefits of combining information-theoretic analyses of experimental data and computational modelling, as integrating datasets from patients with those from healthy subjects. We hope our findings will inspire further work deepening our understanding about the relationship between neural dynamics and high-level brain functions, which in turn may accelerate the development of novel, mechanism-based treatments to foster and promote mental health.

## Acknowledgements

H.R. is supported by the Imperial College President’s PhD Scholarship. P.M. and D.B. are funded by the Wellcome Trust (grant no. 210920/Z/18/Z). F.R. is supported by the Ad Astra Chandaria foundation. C.T. is funded by the Psychedelic Research Group, Imperial College London. R.C.-H. is the Ralph Metzner Chair of the Psychedelic Division, Neuroscape at University of California San Francisco. A.K.S. is supported by the European Research Council (Advanced Investigator Grant 101019254.) This work was supported in part by grant MR/N0137941/1 for the GW4 BIOMED MRC DTP, awarded to the Universities of Bath, Bristol, Cardiff and Exeter from the Medical Research Council (MRC)/UKRI (S.B.). The LSD research described here was supported by the Beckley Foundation and Walacea Crowd Funding campaign. We thank Imperial College London’s Research Computing Service for the computing facilities. Parts of this work were carried out using the computational facilities of the Advanced Computing Research Centre, University of Bristol.

## Appendix A. Further details on the predictive processing model

This appendix outlines how a process of Bayesian inference on the probability distribution described by Eq. (3) can be interpreted in terms of the joint dynamics of prediction and prediction error. This is covered in standard textbooks of time series analysis (e.g. in Ref. [55]) – however, it is provided here for completeness and accessibility.

As a starting point, we assume that a given brain region is trying to infer the hidden cause *h*_*t*_ of its afferent signal, *s*_*t*_. The brain can use all the previous signals, ***s***^*t*−1^ = (*s*_1_, … , *s*_*t*−1_), to generate an optimal prior estimation of *h*_*t*_ , which is given by *p*(*h*_*t*_ | ***s***^*t*−1^).When a new sample *s*_*t*_ is observed, this prior can be updated using Bayes’ rule,

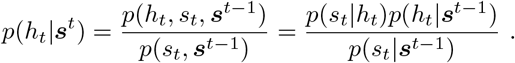

Note that, while in general computing *p*(*h*_*t*_ | ***s***^*t*^) can be computationally challenging, when all the distributions in the right-hand side of the equation above are Gaussian (as per Eq. (3)) the posterior is easily calculable – as we explain below.

For consistency with the model in Eq. (2), we assume that *h*_*t*_ | ***s***^t−1^ is a Gaussian random variable with mean *ĥ*_*t*_ and variance 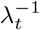. By considering *p*(*h*_*t*_ | ***s***^*t*−1^) as a prior and *p*(*s*_*t*_ | *h*_*t*_) as a likelihood, we can compute the posterior *p*(*h*_*t*_ | ***s***^*t*^) by using Bayes’ rule above for Gaussian variables. In this case, standard results for conditional Gaussian distributions (e.g. Ref. [55, Eq. (4.2)]) show that

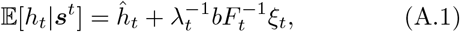

where 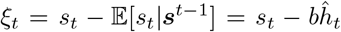 is the error in the prediction of *s*_*t*_ given ***s***^*t*−1^, and 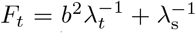 is the predictive covariance of *s*_*t*_ given ***s***^*t*−1^. Then, by using Eq. (3a), one can propagate the prediction in Eq. (A.1) to the next step, and obtain a recurrent update equation for *ĥ*_*t*_ given by

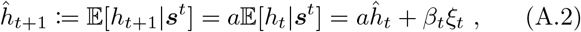

where 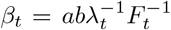 is known as the *Kalman gain* parameter [55, Sec. 4.3] and which, as described in Sec. 2.4, depends only on its previous value *ĥ*_*t*−1_ and the prediction error *ξ*_*t*_.

Furthermore, from the definition of *β*_*t*_ and *F*_*t*_ it can be seen that increasing *λ*_s_ leads to a higher *β*_*t*_, thus increasing the bottom-up influence of prediction errors. A similar argument can be made for the increase of *β*_*t*_ with lower *λ*_p_, although this requires writing a recurrent expression analogous to Eq. (A.2) for *λ*_*t*_ and is more mathematically involved. Interested readers are referred to Sec. 4.3 of Ref. [55] for a detailed derivation.

## Appendix B. Table with definition of the selected ROIs

**Table B.1:**
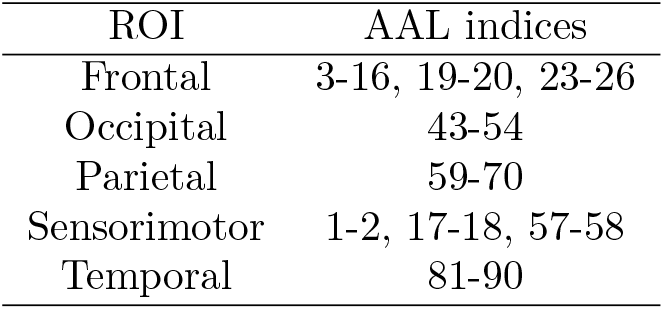
Selected regions from the AAL atlas and their corresponding region of interest (ROI).

## Footnotes

2 Note that in the simple scenario of Eqs. (3), with a single level and Gaussian distributions, inference is in fact exact. In larger or more complicated models inference is often carried out only approximately [47].

